# Nanomechanics of cell-derived matrices as a functional read-out in Collagen VI-related Congenital Muscular Dystrophies

**DOI:** 10.1101/2024.09.13.612824

**Authors:** Tom White, Arístides López-Márquez, Carmen Badosa, Cecilia Jiménez-Mallebrera, Josep Samitier, Marina I. Giannotti, Anna Lagunas

**Affiliations:** Nanobioengineering Group, Institute for Bioengineering of Catalonia (IBEC), The Barcelona Institute of Science and Technology (BIST), Baldiri Reixac, 10-12, 08028 Barcelona, Spain; Laboratorio de Investigación Aplicada en Enfermedades Neuromusculares, Institut de Recerca Sant Joan de Déu, C. de Sta. Rosa, 39, 08950 Barcelona, Spain; Unidad de Patología Neuromuscular, Servicio de Neuropediatría, Hospital Sant Joan de Déu, Pg. de Sant Joan de Déu, 2, 08950 Esplugues de Llobregat, Barcelona, Spain; Centro de Investigaciones Biomédicas en Red de Enfermedades Raras (CIBERER), C. de Melchor Fernández Almagro, 3, Fuencarral-El Pardo, 28029 Madrid, Spain; Department of Genetics, Microbiology and Statistics, Faculty of Biology, University of Barcelona, Diagonal, 643, 08028 Barcelona, Spain; CIBER-BBN, ISCIII, Barcelona, Spain; Department of Electronic and Biomedical Engineering, University of Barcelona, Martí i Franquès, 1, 08028 Barcelona, Spain; Nanoprobes & Nanoswitches Group, Institute for Bioengineering of Catalonia (IBEC), The Barcelona Institute of Science and Technology (BIST), Baldiri Reixac, 10-12, 08028 Barcelona, Spain; Department of Materials Science and Physical Chemistry, University of Barcelona, Martí i Franquès, 1, 08028 Barcelona, Spain

**Keywords:** atomic force microscopy-based force spectroscopy, collagen-VI related congenital muscular dystrophies, extracellular matrix, cell-derived matrices, elastic modulus, gene editing

## Abstract

Atomic force microscopy-force spectroscopy (AFM-FS) applied to biological samples can provide information on the micro- and nanoscale mechanical properties of tissues, which are a determinant of cellular behavior. Mechanical properties are largely determined by the structure and composition of the extracellular matrix (ECM), for which several *in vitro* models have been developed, including cell-derived matrices (CDMs). CDMs are decellularized natural ECMs assembled by cells, that closely mimic the *in vivo* stromal fiber organization and molecular content.

Here we applied AFM-FS to evaluate the nanomechanical properties of CDMs obtained from primary skin fibroblast cultures of patients affected by collagen VI-related congenital muscular dystrophies (COL6-RDs). COL6-RDs are a set of neuromuscular conditions caused by pathogenic variants in any of the three major COL6 genes, which result in deficiency or dysfunction of the COL6 incorporated into the ECM of connective tissues. Current diagnosis includes the genetic confirmation of the disease and categorization of the phenotype based on maximum motor ability, as no direct correlation exists between genotype and phenotype of COL6-RDs.

The development of new tools able to identify phenotype traits can significantly contribute to the diagnosis and prognosis of COL6-RDs. We describe differences in the elastic modulus (*E*) among CDMs deriving from patients with different clinical phenotypes, as well as the restoration of *E* in CDMs obtained from genetically edited cells. Results anticipate the potential of the nanomechanical analysis of CDMs as a complementary clinical tool, providing phenotypic information about COL6-RDs and their response to gene therapies.

## Introduction

The extracellular matrix (ECM) is a three-dimensional (3D) and highly dynamic acellular component present in all tissues. It is formed by a variety of compounds including minerals and a mesh of macromolecules that are produced intracellularly by resident cells. While the ECM provides physical support and controls tissue homeostasis, its dysregulation results in a number of human conditions [1]. In particular, changes in the mechanical properties of the ECM are a hallmark of severe diseases such as pulmonary fibrosis and cancer [2,3], for which a chronic stiffening has been reported and associated with increased secretion of collagens and collagen cross-linkers [4].

Given the relevance of the ECM in disease progression, several *in vitro* models have been developed to study its biochemical and mechanical properties, ranging from the use of decellularized native tissues to the engineering of ECM analogues of different complexity. Decellularized tissue slices can provide a faithful mimic of tissue-specific architecture, yet they cannot be bioengineered. Alternatively, cell-derived matrices (CDMs), produced *in vitro* from patients’ cells, closely mimic the *in vivo* stromal fiber organization and molecular content of the ECM [5–8], as well as allowing for their controlled manipulation at different stages to produce tunable outcomes in matrix production [9]. In CDMs, cells can be genetically engineered to tailor their properties or they can be cultured on substrates with different topographies that are then transferred to the final matrices [10]. In this line, our group has previously published on the production of engineered anisotropic CDMs to study cell growth and directed 3D cell migration [11,12]. More recently, we produced CDMs from patients with collagen type VI-related congenital muscular dystrophies (COL6-RDs) to study the morphology and organization of protein fibers in the ECM. In agreement with previous results obtained in Collagen Type VI-deficient CDMs from stable *COL6A1* knockdown cell lines [13], we found that CDMs from COL6-RD patients presented alterations in structure and composition, showing a significant decrease in COL6 secretion, also affecting the organization of matrix proteins fibronectin and fibrillin-1 [14].

COL6-RDs comprise a heterogenous group of incurable and rare disorders that present different severities of clinical manifestation, spanning from the milder Bethlem myopathy (BM) to the severe Ullrich congenital muscular dystrophy (UCMD), with a range of intermediate cases [15–17]. Underlying all these forms are mutations in any of the three major collagen VI (COL6) genes, *COL6A1, COL6A2, COL6A3* [18]. These genes encode three α chains, α1(VI), α2(VI) and α3(VI) respectively, which assemble first to form a triple-helix monomer before further association to dimers and then tetramers via disulfide bond formation. These tetramers are finally secreted in the ECM producing an extended network of beaded microfilaments [19]. In COL6-RDs, more than 200 mutations in COL6 genes have been identified, predominantly affecting *COL6A1* and *COL6A2*. Exon skips or point mutations proximal to the N-terminal region of the triple helix domain that still allow tetramer assembly and secretion result in the incorporation of aberrant COL6 into the ECM. Conversely, those mutations that lead to very low to no expression of the affected α-chains or that prevent tetramer formation, result in COL6 being almost absent in patients’ muscle and fibroblast cultures [18,20,21]. Current COL6-RD diagnosis includes the genetic confirmation of the disease and categorization of the phenotype based on maximum motor ability [22], as correlations between genotype and phenotype are still very difficult to identify [20]. Therefore, new tools able to recognize phenotype traits can significantly contribute to the diagnosis and prognosis of COL6-RDs.

COL6 was found fundamental in defining a healthy ECM with optimal stiffness. Stress-strain tests conducted in muscle tissue of a total knockout mouse model (*Col6a1*^-/-^) resulted in a significant decrease in stiffness compared to the WT counterparts, which was attributed to the lack of COL6 in the ECM, compromising the *in vitro* and *in vivo* activity of satellite cells and affecting muscle regeneration [23]. Based on these previous results and taking advantage of the reductionist model provided by the CDMs, in the present work we use atomic force microscopy-force spectroscopy (AFM-FS) to evaluate the stiffness in CDMs produced from patients diagnosed with COL6-RDs. We further investigated the effects of gene editing on the mechanical and morphological properties of CDMs, demonstrating the potential of mechanical analysis of patient CDMs as a functional biomarker of COL6-RD progression and therapeutic response.

## Experimental

### Human Fibroblasts

Forearm dermal fibroblasts were used in this study [24]. Dermal fibroblasts were obtained from the Hospital Sant Joan de Deu biobank for five representative COL6-RD patients covering a spectrum of mutations and phenotypes, and two controls unaffected by any neuromuscular condition (Table 1). Primary cells were collected following a standard skin biopsy procedure. All patients had a genetically or pathologically confirmed diagnosis of COL6-RD, with their phenotypes subsequently categorized based on maximum motor ability [22]. Written informed consent was obtained from individuals and/or their parents or guardians. Biological samples were stored and managed by the Hospital Sant Joan de Déu (HSJD) Biobank.

**Table 1.** Description of fibroblast donors included in this study with their relevant genetic background.

| Donor | Phenotype | Mutation | Gene<br>(pattern of inheritance) | Sex | Age at time of biopsy<br>(years) |
| --- | --- | --- | --- | --- | --- |
| Patient 1 | BM | c.877G > A (Gly293Arg) Exon 10 | *COL6A1* (AD) | Female | 9 |
| Patient 2 | Intermediate | c.877G > A (Gly293Arg) Exon 10 | *COL6A1* (AD) | Male | 8 |
| Patient 3 | Intermediate | c.2329T > C (Cys777Arg) | *COL6A2* (AD) | Female | 12 |
| Patient 4 | UCMD | c.901-2A > G Intron 7 | *COL6A2* (AD) | Female | 3 |
| Patient 5 | UCMD | c.930 + 189C > T Intron 11 | *COL6A1* (AD) | Male | 4 |
| Control 1 | - | - | - | Male | 1 |
| Control 2 | - | - | - | Male | 2 |

The present study was performed in accordance with the Declaration of Helsinki. All experimental protocols were approved by the Fundació Sant Joan de Déu Ethics Committee on Clinical Research (project identification code: PIC-24-23). Methods were carried out in accordance with relevant guidelines and regulations.

### Genetically edited cells

Genetically edited cells from López-Márquez *et al*. [25] were used. In summary, CRISPR/*Cas*9-based genome editing was performed on dermal fibroblasts of two patients presenting either the BM (patient 1) or intermediate phenotype (patient 2). Two different CRISPR RNA guides, crRNA_1 (nucleotide sequence: *5’-CCTGGTACCCAACAGGTCTG-3’)* and crRNA_2 (nucleotide sequence: *5’-CCCGGGGACCTCAGACCTGT-3’)*, were used to silence the dominant-negative *COL6A1* Het. c.877G>A; p.Gly293Arg pathogenic variant present in both individuals. Dermal fibroblast cultures were transfected with the ribonucleoprotein complex formed by the *Cas* 9 endonuclease and the duplex RNA from the union of each of the crRNAs and a fluorescently labeled transactivating crRNA (tracrRNA-ATTO550). Transfection efficiency was measured by fluorescence-activated cell sorting (FACS) using the tracrRNA-ATTO550 fluorophore, resulting in >80% of ATTO550 positive cells for any of the two guides used. Editing success was evaluated by analyzing the DNA reads from transfected cells using Mosaic-Finder [26]. WT *COL6A1* allele resulted almost unaltered, while the mutant allele was reduced more than 50%, demonstrating the specificity of the process. Allele specific expression of *COL6A1* was analyzed by ddPCR using specific probes for WT or mutant cDNA. The normalized expression of the mutant allele was reduced, reaching >80% in Patient 1 treated with crRNA_2 (BM_Edited2). Please refer to the paper from López-Márquez *et al*. [25] for details.

### Cell culture

Donor fibroblasts were maintained at 37°C and 5% CO_2_ in a high glucose Dulbecco’s modified Eagle’s Medium (DMEM) (Invitrogen) supplemented with 10% fetal bovine serum (FBS) (Sigma-Aldrich), 1% penicillin and streptomycin (Invitrogen) and 1% sodium pyruvate (Sigma-Aldrich). The growth medium was replaced every 2 days until 80% confluence was reached at which point cells were harvested. To achieve detachment, cells were incubated with 0.25% trypsin-EDTA (Life Technologies) at 37°C and 5% CO_2_ for 5 minutes. Supplemented growth medium was then added to inactivate trypsin, and the cell suspension was transferred to a 15 ml tube for centrifugation at 500 rpm for 5 min. The supernatant was discarded, and the pellet resuspended in 10 ml of supplemented growth medium. Using a Neubauer chamber, the number of cells and thus the concentration per ml of suspension could then be calculated for subsequent CDM deposition.

### CDM production

CDMs were generated as previously described [11,12,14]. Precleaned glass microscope coverslips (Ø 18 mm, Superior Marienfeld) were incubated with a sterile 1% gelatin (from porcine skin Sigma-Aldrich) solution in PBS (Sigma-Aldrich) at 37°C. A crosslinking reaction for the gelatin layer was initiated with 1% filtered glutaraldehyde in PBS (Sigma-Aldrich) and then quenched with 1 M sterile glycine (Sigma-Aldrich) solution in PBS. The coated coverslips were stored in the fridge with 1% penicillin and streptomycin in PBS for a maximum of two weeks until use.

To produce the CDMs, growth medium containing 4 × 10^4^ cells/cm^2^ of primary fibroblasts was added to the coated substrate. After the fibroblasts had formed a confluent monolayer, 50 μg/ml ascorbic acid (AA) in PBS (Sigma-Aldrich) was added every day to stimulate collagen production. Cell media was replaced every three days. After 7 days of AA treatment, samples were decellularized with a filtered extraction buffer: 0.5% Triton X-100 (Sigma-Aldrich) and 300 mM of ammonium hydroxide solution (NH_4_OH, Sigma-Aldrich) in PBS. After removal of cell media, this solution was added gently on top of the samples for 4 min before being diluted in 2 ml of PBS. Cell detachment can be monitored under light microscopy [27]. Matrices were stored overnight at 4°C and then washed carefully multiple times to remove any cell debris. To evaluate the effectiveness of the decellularization process, the absence of cellular structures such as cell nuclei was confirmed by IF, as previously described [14].

### Immunostaining

CDMs were fixed with formalin 10% (Sigma-Aldrich) for 20 min, before further treatment with ammonium chloride 50 mM (Sigma-Aldrich) in PBS for 20 min to quench any free aldehyde groups. Blocking and permeabilization was then carried out with a solution of 1% bovine serum (BSA) (Sigma-Aldrich) and 0.1% saponin (Sigma-Aldrich) in PBS for 10 min. The matrices were next incubated with the primary antibody mouse monoclonal anti-Collagen VI antibody (Millipore) (1:400) in a PBS solution with 1% BSA and 0.1% saponin for 1 h at room temperature. Samples were carefully rinsed before addition of the fluorophore conjugated secondary antibody: Alexa 488 goat anti-mouse (Thermo Fisher Scientific) (1:1000) in PBS with 1% BSA and 0.1% saponin for a further 1 h at room temperature. Staining with Hoechst 33342 (Molecular Probes) (1:1000) was also performed to detect any nuclear remains as a verification of successful decellularization. Samples were finally rinsed with PBS and then mounted with Fluoromount (Sigma-Aldrich) on 76×26 mm microscope slides (RS FRANCE). The samples were allowed to set overnight at room temperature before imaging.

### Confocal microscopy

The Zeiss LSM 800 (Carl Zeiss Microscopy GmbH, Jena, Germany) confocal laser scanning microscope with a 20x air (0.8 NA) Plan-Apochromat objective lens was used to acquire 16-bit IF images. Diode lasers with wavelengths 405 and 488 nm were used to excite the relevant fluorophores with emission detection bandwidths configured at 400–500 nm (Hoechst 33342) and 400–570 nm (fluorochrome Alexa 488). Z-stacks were acquired in increments of 1 μm for an average of 20 to reconstruct the 3-dimensionality of the CDM. In total COL6 data was acquired for a minimum of 3 fields of view per replicate with at least two replicates per donor. Laser and detector settings were maintained constant across all samples. Confocal stacks were visualized as maximum intensity projections using Fiji software [28]. CurveletTransform—Fiber Extraction (CT-FIRE) algorithm was then used to segment individual protein fibers and calculate biophysical parameters, as previously described [14].

### AFM-FS

AFM-FS experiments were performed on the decellularized CDMs, at room temperature under liquid environment (PBS buffer solution pH 7.4, 10 mM phosphate, 2.7 mM KCl and 137 mM NaCl), following a well-established method to quantify the elastic properties of ECM models, including decellularized tissue ECM [29]. AFM-FS experiments were done with an MFP-3D atomic force microscope (Asylum Research) using a V-shaped Si_3_N_4_ cantilever with a 10 μm diameter, polystyrene colloidal probe and a nominal spring constant of 0.06 N/m (NOVASCAN). The sensitivity of the photodiode was acquired by recording a deflection-Z position curve on a mica substrate before the cantilever spring constant was calibrated according to the thermal oscillation technique [30]. AFM-FS measurements were performed by approaching and retracting the colloidal probe to the sample at a constant velocity of 3 μm/s, to a maximum applied force around 600 pN. Force-indentation curves were recorded in force map mode acquiring 36 curves within a total scan size of 60 × 60 μm^2^. A minimum of 5 distinct sample regions were selected per sample for the force mapping with at least two replicates per donor measured. Measurements were performed on all donors except patient 5 and control 2 (Table 1). The Young’s elastic modulus *E* value was determined from these curves by fitting the Hertz model using IGOR Pro 6.22A software to indentation curves (Fig. S1), using a Poisson ratio ν = 0.5, without the need of any correction factor. The fitting was done to a maximum force bellow 400 pN, which ensured an indentation of less than 15% of the sample thickness, estimated to be around 6-10 μm (Fig. S2) [14], thus avoiding any mechanical influence from the underlying substratum [29,31–33]. Curves were individually analyzed and discarded when excessive adhesion between the sample and tip prevented a good quality of fit for the model.

### Statistical Analysis

All statistical analysis and plotting of quantitative data were performed with GraphPad Prism 9. Normality tests were performed before any statistical comparison. For normal distributions, significant differences were assessed with the two-tailed Student’s t-test or Tukey’s multiple comparisons test for comparison of more than two groups. When data was significantly drawn from a normal distribution, significant differences were assessed with the Mann-Whitney test or Kruskal-Wallis test for comparison of more than two groups. Statistical significance was assumed for α < 0.05. At least three replicates per condition and 2 independent experiments were performed in each case (N = 2). When results are grouped together into a single phenotype obtained from more than one patient it is because there are no significant differences among the individual patients.

## Results

### Mechanical analysis of CDMs from COL6-RDs patients

In the present study, human dermal fibroblasts were used. Although producing inferior levels of ECM when compared to fibro/adipogenic progenitor cells (FAPs), which are the main ECM source [34–36], dermal fibroblasts are easily accessible, expandable cells in culture that express high levels of the 3 major COL6 genes [37].

Human fibroblasts were isolated from forearm skin biopsies of five representative patients covering a spectrum of mutations and phenotypes, and two healthy donors. Their COL6-RD phenotypic classification and genetic background is summarized in Table 1. CMDs were produced as described in previous publications from our group [11,12,14]. Briefly, confluent cultures of plated fibroblasts were treated with ascorbic acid (AA) for 7 consecutive days stimulating the secretion of collagen. Cellular components were subsequently removed, leaving a 3-dimensional scaffold representative of individual COL6-RD cases.

To evaluate the mechanical properties of the CDMs of the different patients (and phenotypes), we measured the CDM’s resistance to deformation and calculated the Young’s elastic modulus, *E* using AFM-FS. AFM-FS is a suitable technique to evaluate the mechanical characteristics of ECM models at the micro- and nanometer scale, where mechanical interactions between cell membrane receptors and matrix ligands take place [29].

Owing to the heterogenous microenvironment of CDM models, high variability within the same sample can occur for the mechanical response at the nanometer scale. Thus, we selected a spherical geometry (diameter = 10 μm) for the AFM tip to maximize tip-surface contact area and enable the mechanical contribution of multiple matrix fibers to be averaged in a single measurement. From this, we obtain an effective mesoscopic elasticity to compare different samples and conditions, that is more relevant to clinics and diagnostics than that of individual nanoscale components [29,38–40].

Additionally, selecting a spherical tip is advisable for very soft samples, to generate small strains [41]. Force-indentation curves of the different COL6-RDs CDMs displayed clear differences with respect to the contact region response and thus deformability of the matrix (representative curves are shown in Fig. 1a). Before quantification of the CDM’s elasticity, the curves were assessed individually to ensure excessive adhesion between the probe and sample was not present. Interestingly, this phenomenon was encountered more frequently for UCMD samples, with many pulling events observed in the retraction curves (Fig. 1a). These are typical patterns for polymeric mesh/network pulling. The Hertz contact model was subsequently applied to the appropriate indentation curves, using *E* as a fitting parameter (Fig. S1). We found the *E* value for CDM from healthy donors (control) to be 1736 ± 100 Pa, with COL6-RD patients CDMs displaying significant deviations from this ‘healthy’ range (Fig. 1b). Whilst an increase in stiffness with *E* of 3313 ± 200 Pa (mean ± SEM) was observed for the intermediate phenotype, CDMs *E* value was reduced to 659 ± 50 Pa and 152 ± 6 Pa correspondingly for BM and UCMD patients. *E* for Intermediate patients 2 and 3 were not significantly different, with values of 3336 ± 200 Pa and 3019 ± 300 Pa, respectively. Interestingly, a five-fold increase in *E* was calculated for intermediate patient 2 over BM patient 1 CDMs, despite both sharing the same mutant variant. The more frequent observation of pulling events in the BM and, especially, UCMD samples along retraction of the AFM probe (Fig. 1a) suggests a less tight mesh, making it easier to separate and stretch fibers out the ECM. Therefore, the mechanical properties of the CDMs form COL6-RD patients are significantly different from those from healthy donors, but also among the different phenotypes.

**Fig. 1.**
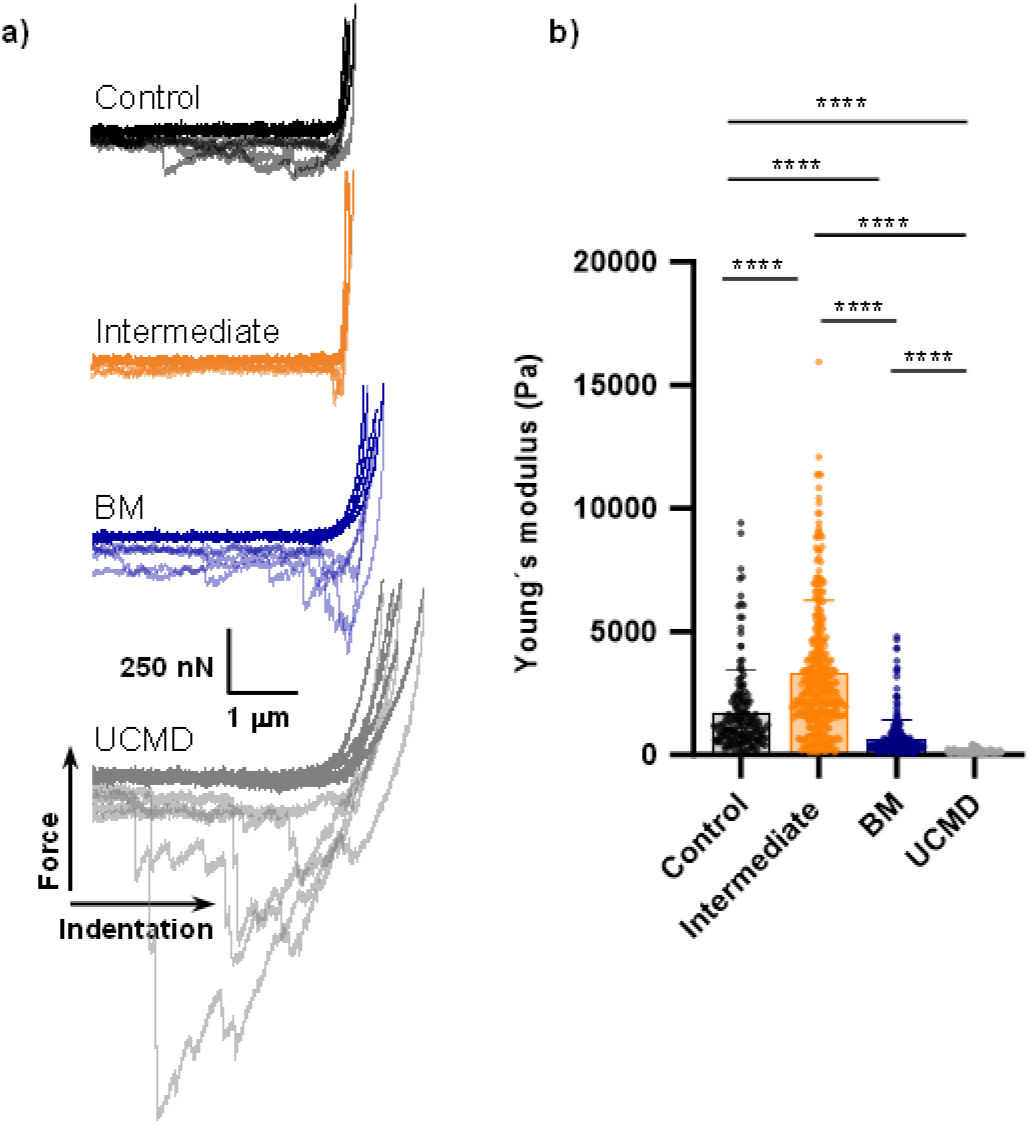
Mechanical analysis of CDMs. **a)** Representative force-indentation curves for each condition tested (dark: approach, light: retraction). **b)** Values of the Young’s modulus € for CDMs obtained from healthy donors (control) and COL6 RD patient cells: Intermediate (patients 2 and 3), BM (patient 1) and UCMD (patient 4) phenotypes. Results (bars graphs) are the mean ± SD. ****p < 0.0001.

### CDMs derived from patient fibroblasts genetically edited

Genetically edited cells from López-Márquez *et al*. [25] were used. Dermal fibroblasts from patients bearing the pathogenic variant *COL6A1* Het. C.877G>A, presenting either the BM (patient 1, Table 1) or intermediate (patient 2, Table 1) phenotype were edited by CRISPR/*Cas*9-based gene editing. Two different CRISPR RNA guides, crRNA_1 and crRNA_2, were used to silence the dominant-negative *COL6A1* c.877G>A; p.Gly293Arg pathogenic variant. The effectiveness of the editing strategy was analyzed at the genomic DNA level by Next Generation Sequencing, observing the high specificity of both RNA guides, since the WT allele was not edited at significant levels. Furthermore, most of the allelic variants generated after editing the mutant allele were due to out-of-frame Indels that would predictably lead to specific silencing of the mutant allele [25]. To demonstrate this, specific probes were designed to measure mRNA expression levels of the *COL6A1* mutant and WT allele. Using ddPCR, it was concluded that the reduction in expression of the mutant allele was at least 60%, being 80% in the case of patient 1treated with crRNA_2. The WT allele was expressed at normal levels compared with the unedited fibroblasts [25]. To examine the response of patient’s fibroblasts (patient 1) and (patient 2) to the gene silencing of their mutant allele, CDMs were produced and analyzed by immunofluorescence and image processing, and by AFM-FS.

#### Immunofluorescence analysis of CDMs derived from treated patient fibroblasts

The secreted COL6 was stained in the CDMs. Representative confocal z-projections obtained for COL6 staining in CDMs from control and COL6-RD samples, produced by edited and unedited cells, are shown in Fig. 2a. Visual inspection of the images shows fewer and more segmented fibers for patients with COL6-RDs compared to the control sample, as well as some degree of morphological recovery in the edited samples, especially for the intermediate phenotype. Accordingly, the results from image processing (Fig. 2b) show that fewer, shorter and thinner COL6 microfilaments are produced in patient samples compared to controls, and that they are significantly less aligned for the intermediate phenotype. Gene editing in general for the intermediate phenotype produces a leveling of the evaluated parameters toward control values, leading to a significant increase in the number, length and width of COL6 microfilaments present in the CDMs, especially when using the crRNA_2 guide (Edited 2). However, COL6 control levels are not fully recovered, as expected since *COL6A1* allele is partially silenced.

**Fig. 2.**
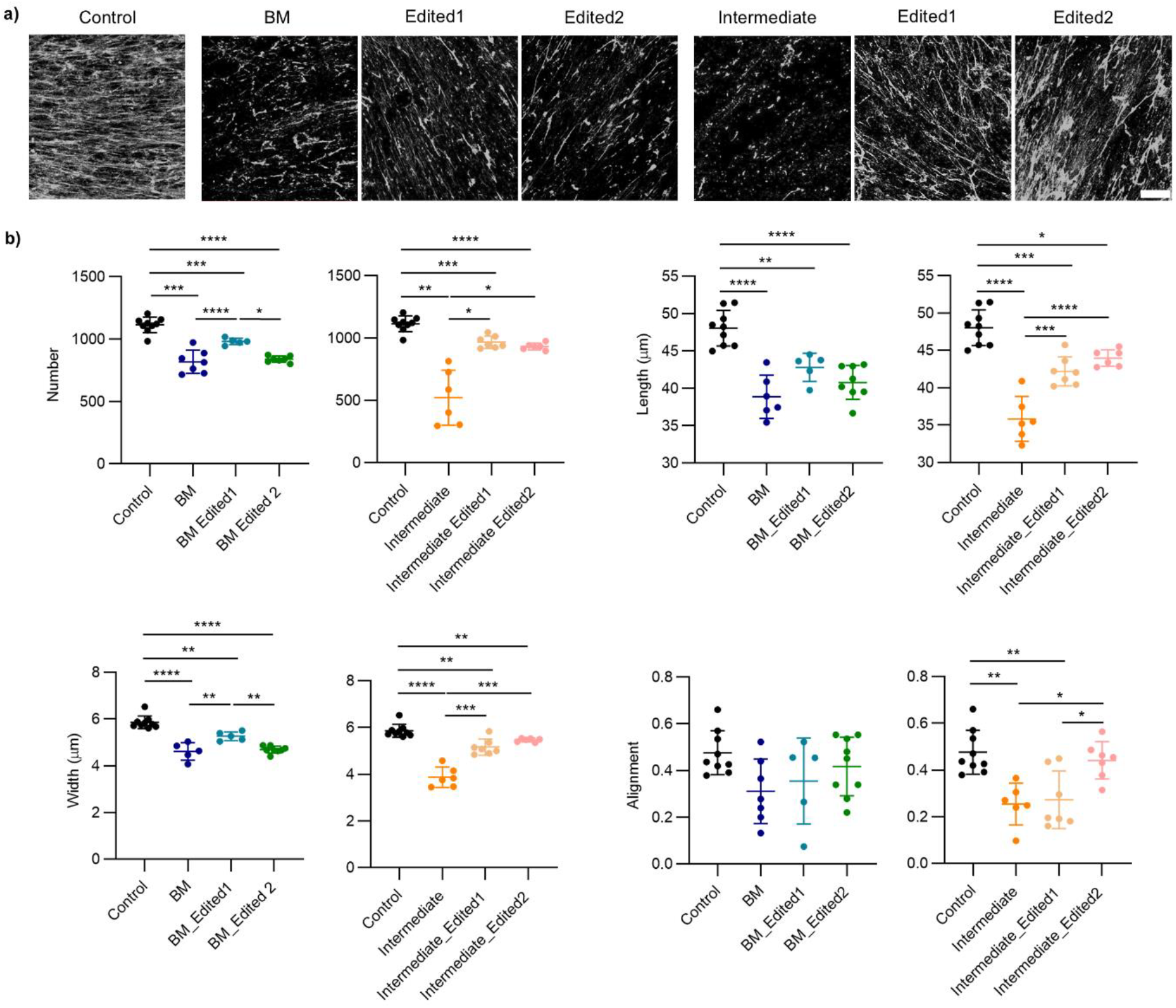
Effect of gene editing on COL6 organization in CDMs. **a)** Representative confocal z-projections of immunostained COL6 in the CDMs: from fibroblasts of healthy donors (Control) and from COL6-RD patients 1 and 2 with BM and intermediate phenotypes, respectively, before and after gene editing using either crRNA_1 (Edited1) or crRNA_2 (Edited2). Scale bar = 50 μm. **b)** Graphs displaying various characteristics of COL6 microfilaments obtained by image processing using CT-FIRE algorithm. Results are the mean ± SD. *p < 0.05, **p < 0.01, ***p < 0.001 ****p < 0.0001.

Strikingly, while editing with the crRNA_2 guide statistically levels control values for microfilament alignment, the use of the crRNA_1 guide (Edited 1) does not produce significant differences with respect to the unedited sample. In this regard, López-Márquez et al. also reported variations in patient response as a function of the RNA guide used, with crRNA_2 often recovering more COL6 signal [25].

Alterations in COL6 arrangement after gene editing were less evident for the BM phenotype. In this case, there was a similar trend towards levelling control values in samples edited with cRNA_1, although the increase was only statistically significant for the number and width of secreted COL6 microfilaments.

#### Mechanical analysis of CDMs derived from treated patient fibroblasts

Markedly, matrices derived from gene-edited cells all obtained values of *E* comparable to the healthy donor (control), independently of the RNA guide used (Fig. 3a). For BM_Edited1 and BM_Edited2, we found the value of *E* to be 1420 ± 90 Pa and 1894 ± 80 Pa (mean ± SEM), respectively, displaying a greater than two-fold increase relative to CDMs from unedited cells of BM patient 1. An increase in compliancy of matrices from the non-treated cells of intermediate patient 2 was additionally achieved, with *E* calculated as 1380 ± 70 Pa considering Intermediate_Edited1 and 1593 ± 100 Pa for Intermediate_Edited2. In all cases, no significant difference between edited samples and the control (healthy donor) could be highlighted (Fig. 4a). Figure 4b shows representative force-indentation curves for the edited samples.

**Fig. 3.**
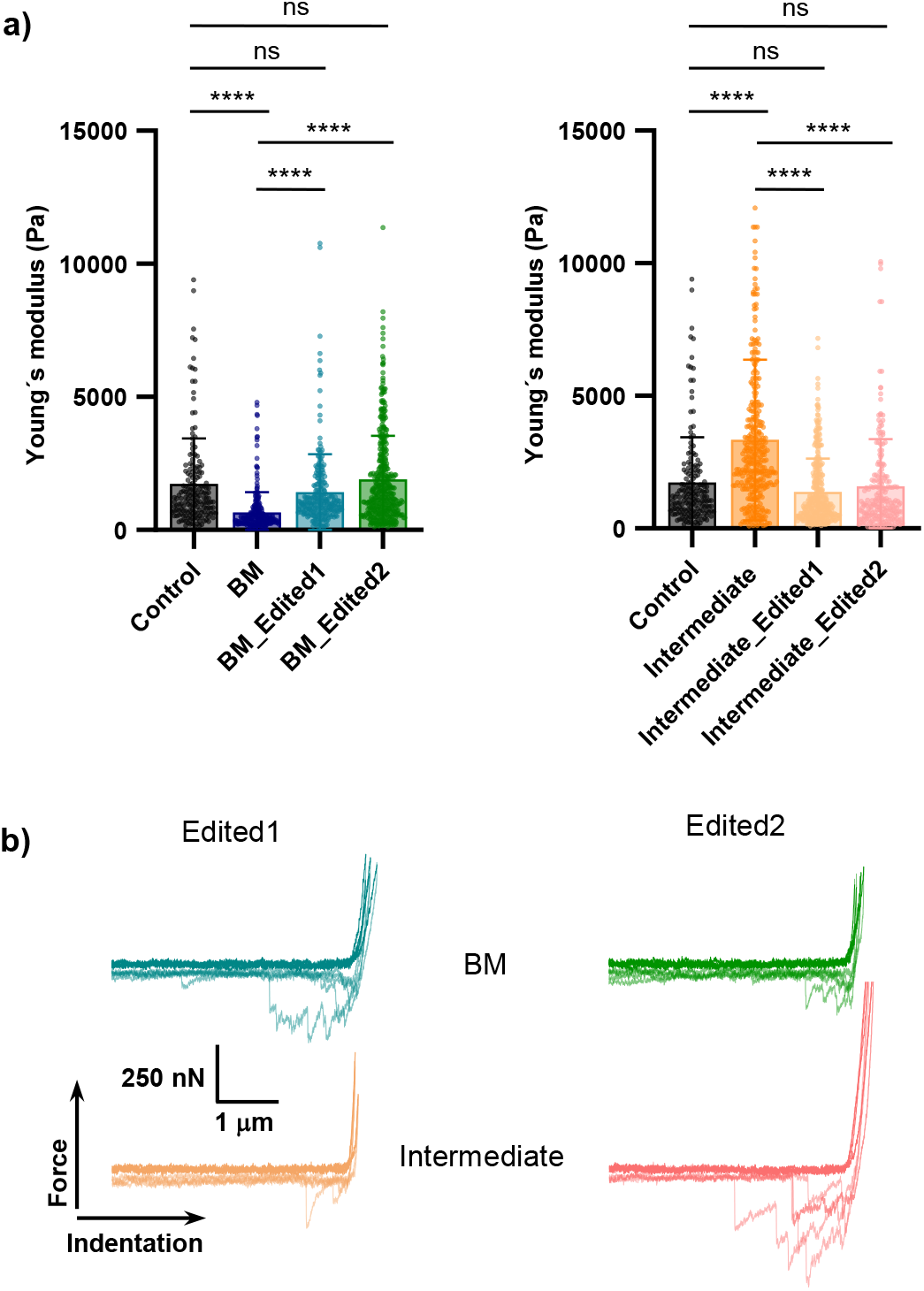
Mechanical analysis of CDMs derived from edited COL6-RD patient fibroblasts. **a)** Values of the Young’s modulus (*E*) for CDMs from fibroblasts of healthy donors (Control) and from COL6-RD patients 1 and 2 with BM and intermediate phenotypes, respectively, before and after gene editing using either crRNA_1 (Edited1) or crRNA_2 (Edited2). Results (bars graphs) are the mean ± SD. ****p < 0.0001, ns = no significant difference. **b)** Representative force-indentation curves for CDMs from COL6-RD patients 1 and 2 with BM and intermediate phenotypes, respectively, using either crRNA_1 (Edited1) or crRNA_2 (Edited2) (dark: approach, light: retraction).

**Fig. 4.**
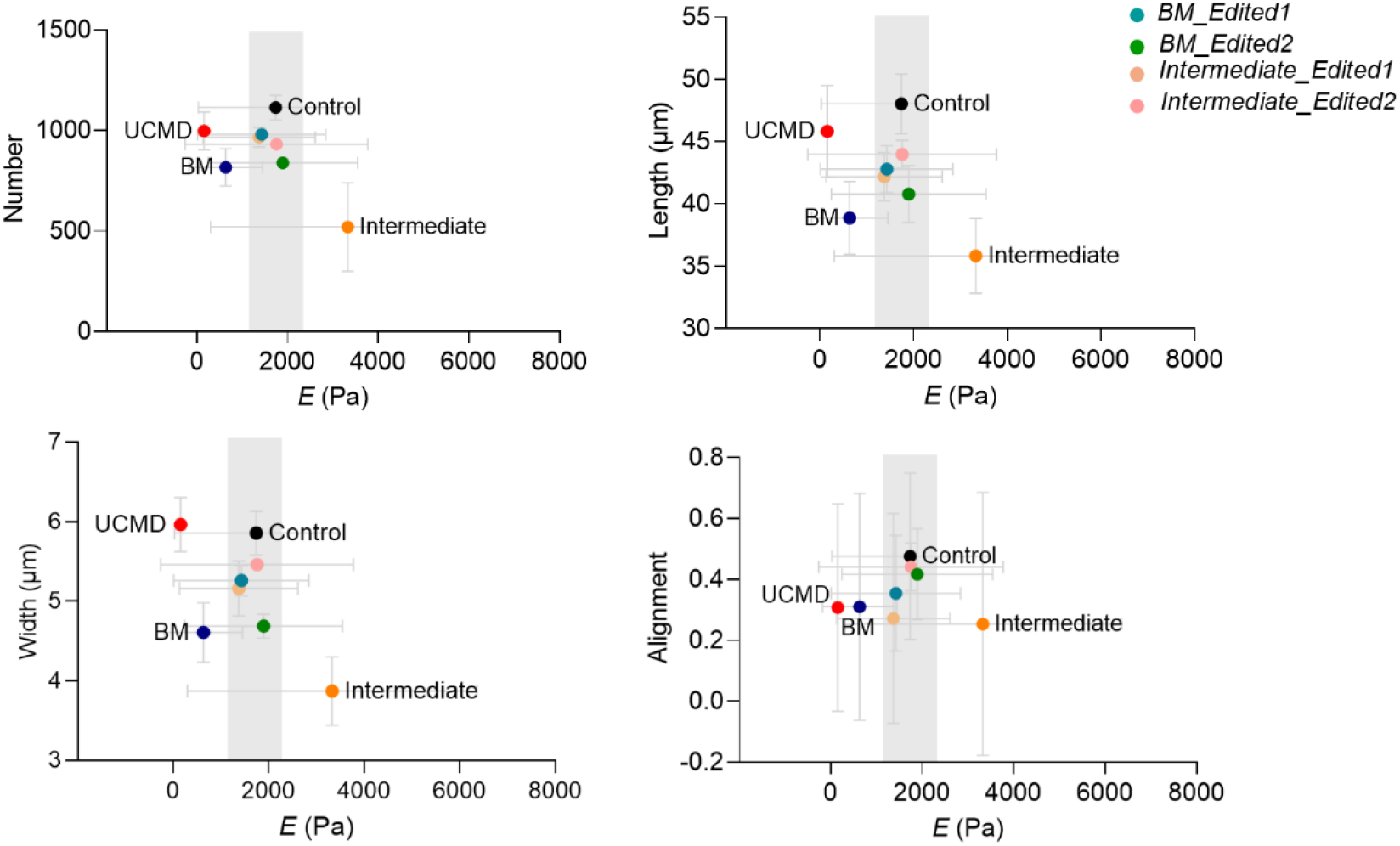
Correlation between morphological and mechanical characteristics. Representation of the COL6 parameters determined by IF and image processing against the CDM’s stiffness (*E*) measured by AFM-FS. The pale grey areas represent the recovery towards control values in the edited samples. Values are the mean ± SD.

Although gene editing does not completely restore COL6, the changes in elasticity of CDMs match that of control samples, thus defining a healthy range for E that excludes the different COL6-RDs phenotypes (estimated as pale grey areas in Fig. 4).

## Discussion

We used AFM-FS to investigate the mechanical properties of CDMs from COL6-RDs patients’ fibroblasts. COL6-RDs represent a group of neuromuscular disorders affecting the ECM of connective tissues. Patients affected by COL6-RDs produce a lower amount of COL6 with altered assembly and interaction with other components of the ECM [24,42]. The ECM’s mechanical response is determined by the organization and relative disposition of its different components [43]. We therefore assessed whether the observed alterations in ECM assembly of COL6-RD patients translate to particular mechanical fingerprints. Recent studies postulate CDMs as a reductionistic *in vitro* model capable of recapitulating the native architecture and composition of the ECM, allowing the identification of the different pathophysiological characteristics of the tissue [9,44]. In the present work we use atomic AFM-FS to obtain the young’s modulus (*E*) as representative of the stiffness in CDMs from patients diagnosed with COL6-RDs.

We found the stiffness of CDMs from COL6-RD patients to significantly deviate from those produced by healthy cells, presenting either an increase or a reduction in *E* depending on the phenotype. A softening of the matrix for BM sample compared to controls was identified. This agrees with previous results conducted in a total knockout mice model (*Col6a1*^*-/-*^), for which a decrease in muscle stiffness was observed [23]. COL6-defficient mice, in general, behave similarly to BM phenotypes [45].

The CDMs from UCMD presented the lowest *E* values. Additionally, pulling events frequently observed in the retraction force curves on UCMD samples suggest a less densely integrated network of fibers. This COL6-RD phenotype is associated with earlier and more severe muscle weakness. Counterintuitively, the Intermediate phenotype does not produce *E* values between the UCMD and BM phenotypes, but results in significantly higher *E* values. In this context, we have previously described that patients with the intermediate phenotype produce significantly straighter and more aligned fibronectin fibers [14], possibly contributing to increased CDM stiffness, as it is known that aligned fibronectin fibres act as a template for the assembly of collagens [46]. Additionally, in studies performed with advanced confocal microscopy on the ECM of the patient with the intermediate phenotype, a significant number of abnormal aggregates positive for collagen VI were observed [25], presumably the product of deficient assembly of the protein. The abundance of these aggregates could also be contributing to an increase in the stiffness of the ECM [47].

Stiffness can produce different mechanical stimuli, which can regulate cell activity both at local and systemic levels through the process of mechanotransduction [48]. As a result, any perturbations to matrix rigidity will generate responses in cells, which could affect their survival, proliferation and migration [49]. Our findings for *E* might therefore help explain impaired processes previously recorded for COL6-RD patients including increased rates in mitochondrial autophagy and apoptosis in skeletal muscle from mice featuring a knockout of the *Col6a1* locus [19], and an accumulation of endosomes-lysosomes and altered mitochondria and Golgi apparatus morphology in primary COL6-RD human fibroblasts [50].

Whilst no effective treatment is currently available for COL6-RDs, novel nucleic acid therapeutics (NATs) have recently emerged as a strategy to target different classes of mutation responsible for the disease [25,37]. Here we take advantage of CDMs combined with AFM-FS to evaluate the effects of a CRISPR system recently developed by López-Márquez *et al*., which was able to silence the mutant allele responsible for substituting glycine 293 for an arginine at the N-terminal in the triple helix of COL6. Two CRISPR RNA guides, namely crRNA_1 and crRNA_2, directed *Cas*9-mediated endonuclease activity to this mutant allele inducing indels, a proportion of which significantly reduced its expression [25]. Matrices derived from gene-edited patient fibroblasts were analyzed in comparison with those produced by the same unedited cells. The immunofluorescence study of secreted COL6 shows that for the intermediate phenotype patient there is a recovery of the values of the different characteristics evaluated, towards the control values, especially when using the crRNA_2 guide (Edited 2). However, although the same trend is intuited for the BM phenotype, significant differences were only found for the number of COL6 microfilaments and their width in the crRNA_1-edited samples (Edited 1). This is in agreement with previous results of López-Márquez et al., achieving higher percentage increases of COL6 intensity in the ECMs of the edited intermediate phenotypes [25].

From AFM-FS studies, we demonstrate how fiber organization affects matrix stiffness and that it is possible to define a healthy range of *E* that excludes different COL6-RD phenotypes. After genetic correction, the stiffness determined for patient matrices shows no significant difference with respect to control (healthy donors) CDMs, suggesting a restoration of ECM mechanical properties, regardless of phenotype or crRNA guide used. This indicates that, although gene editing does not completely restore COL6 levels, the resulting changes are sufficient to significantly improve the mechanical characteristics of the matrix. Thus, analysis of the samples by AFM-FS would provide a more global view of the changes produced by gene editing, highlighting the mechanical characteristics of the matrix as a sensitive biomarker [51] for the evaluation of NATs applied to COL6-RDs.

In conclusion, using CDMs from COL6-RDs patients’ fibroblasts and AFM-FS, it has been possible to establish a healthy range of Young’s modulus, from which the different diseased phenotypes deviate and are distinguishable from each other. We evaluated the response of different patient cells to a recently developed CRISPR/Cas 9 gene correction strategy for which we observed a restoration of matrix stiffness to control values. Thus, variations in the stiffness of CDMs measured with AFM-FS appear to reflect the cascade of events starting from a specific mutation to the final phenotypic expression. The presented results therefore suggest that the mechanical analysis of patients’ CDMs could help either in early stratification of patients, as well as contribute to the evaluation of new therapeutic strategies.

## Credit authorship contribution statement

**A. Lagunas:** Conceptualization, Methodology, Formal analysis, Writing – Original Draft, Visualization, Writing – Review & Editing, Supervision, Project administration, Funding acquisition. **M. I. Giannotti:** Conceptualization, Methodology, Formal analysis, Writing – Original Draft, Visualization, Writing – Review & Editing, Supervision, Project administration, Funding acquisition. **C. Jiménez-Mallebrera:** Methodology, Resources (obtention of patient samples), Supervision, Writing – Review & Editing, Project administration, Funding acquisition. **J. Samitier:** Conceptualization, Writing – Review & Editing, Supervision, Project administration, Funding acquisition. **C. Badosa:** Methodology, Investigation, Writing – Review & Editing. **T. White:** Methodology, Investigation, Formal analysis, Visualization, Writing – Original Draft. **A. López-Márquez:** Methodology, Investigation, Resources (gene editing), Formal analysis, Writing – Review & Editing.

## Data availability

Data will be made available on request.

## Acknowledgements

We acknowledge M. Funes, M. Pérez and E. Almici for technical support and F. S. Tedesco for fruitful discussions. We also acknowledge the Advanced Digital Microscopy Unit at the Institute for Research in Biomedicine (IRB Barcelona). We thank the Nanometric Techniques Unit from the Scientific and Technological Centers of the University of Barcelona (CCiT-UB) for technical assistance. We are grateful to Fundación Noelia for their support to the Jimenez-Mallebrera’s lab at IRSJD and the Biobank HSJD. We acknowledge all the individuals and their families for their support, collaboration, and encouragement. We are indebted to the Biobank of the Hospital Sant Joan de Déu.

Financial support from “Plan Complementario de Biotecnología aplicada a la Salud”, coordinated by Institute for Bioengineering of Catalonia (IBEC) within the framework of “Plan de Recuperación, Transformación y Resiliencia (C17.I1) – financed by the European Commission – NextGenerationEU”. Financial support from Spanish Ministry of Science and Innovation (project PID2022-140459OB-I00 funded by MCIN/AEI/10.13039/501100011033/ and by FEDER A way of making Europe), Department of Research and Universities of the Generalitat de Catalunya (2021 SGR 01545, 2021 SGR 01410), Instituto de Salud Carlos III (ISCIII, grants PI19/0122 and PI22/01382), CERCA Programme / Generalitat de Catalunya, and Networking Biomedical Research Center (CIBER) of Spain. CIBER is an initiative funded by the VI National R&D×i Plan 2008-2011, Iniciativa Ingenio 2010, Consolider Program, CIBER Actions and the Instituto de Salud Carlos III, with the support of the European Regional Development Fund (ERDF).

## Ethical disclosure

The authors state that they have obtained appropriate institutional review board approval and have followed the principles outlined in the Declaration of Helsinki for all human experimental investigations. Written informed consent was obtained from individuals and/or their parents or guardians. Biological samples were stored and managed by the Hospital Sant Joan de Déu (HSJD) Biobank.

---

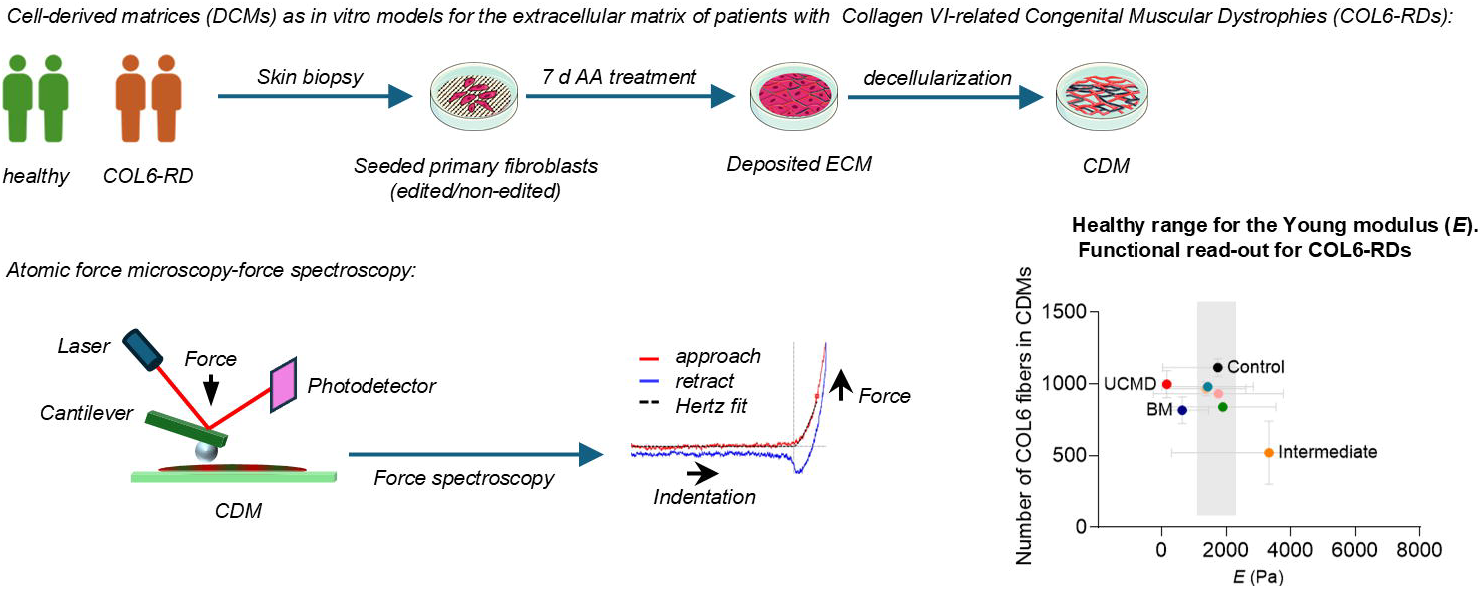

