## Supplementary Information for "Nanomechanics of cell-derived matrices as a functional read-out in Collagen VI-related Congenital Muscular Dystrophies"

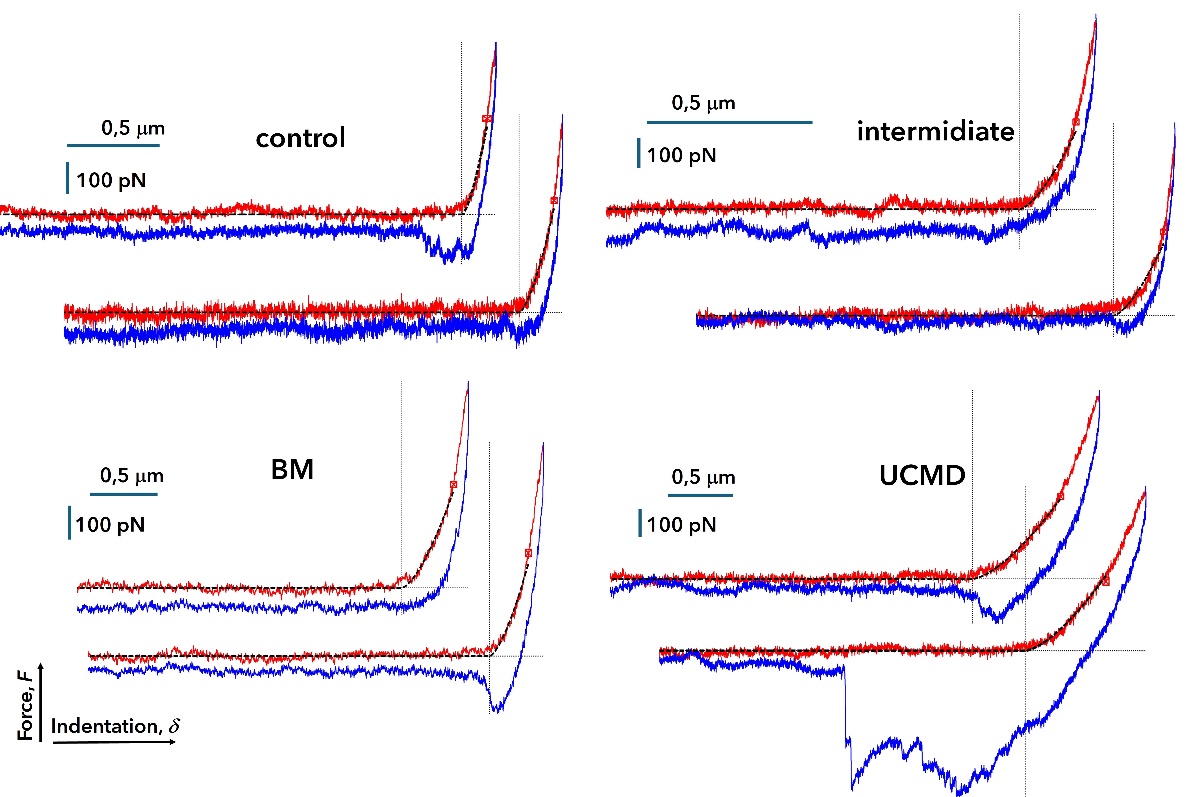

**Figure S1.** AFM-FS force-indentation example curves for CDMs from healthy (control) and COL6-RD (BM, intermediate and UCMD) donors. Approach: red; retract: blue; Hertz fitting (up to red dot): black. Contac point is indicated as the crossing of the dotted vertical and horizontal lines. The Hertz contact model was applied to the approach curves, using *E* as a fitting parameter ($F=\frac{4E}{3(1-v^{2})}R^{2}\delta^{3/2}$).

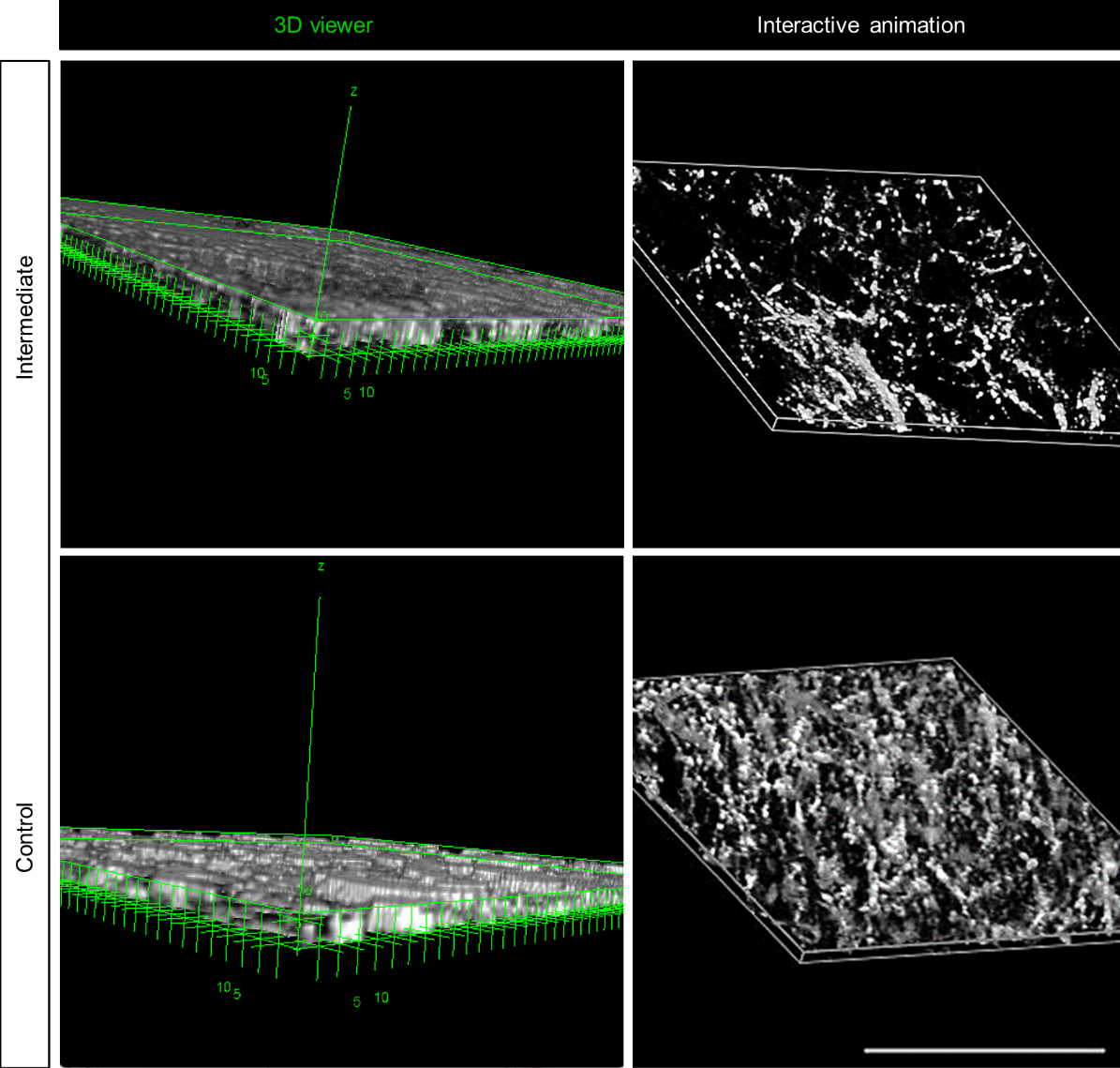

**Figure S2.** Representative examples of thickness estimation in cell-derived matrices (CDMs) through the three-dimensional (3D) reconstruction of confocal images of COL6. Scale bar = 100 µm [1].
